# Comparison of feature representations in MRI-based MCI-to-AD conversion prediction

**DOI:** 10.1101/213132

**Authors:** Marta Gómez-Sancho, Jussi Tohka, Vanessa Gómez-Verdejo, for the Alzheimer’s Disease Neuroimaging Initiative

## Abstract

Alzheimer’s Disease (AD) is a progressive neurological disorder in which the death of brain cells causes memory loss and cognitive decline. The identification of at-risk subjects yet showing no dementia symptoms but who will later convert to AD can be crucial for the effective treatment of AD. For this, Magnetic Resonance Imaging (MRI) is expected to play a crucial role. During recent years, several Machine Learning (ML) approaches to AD-conversion prediction have been proposed using different types of MRI features. However, few studies comparing these different feature representations exist, and the existing ones do not allow to make definite conclusions. We evaluated the performance of various types of MRI features for the conversion prediction: voxel-based features extracted based on voxel-based morphometry, hippocampus volumes, volumes of the entorhinal cortex, and a set of regional volumetric, surface area, and cortical thickness measures across the brain. Regional features consistently yielded the best performance over two classifiers (Support Vector Machines and Regularized Logistic Regression), and two datasets studied. However, the performance difference to other features was not statistically significant. There was a consistent trend of age correction improving the classification performance, but the improvement reached statistical significance only rarely.

## 1. Introduction

Alzheimer’s Disease (AD) is a progressive neurological disorder in which the death of brain cells causes memory loss and cognitive decline. The progression of the neuropathology in AD starts long before clinical symptoms of the disease become apparent [1, 2, 3, 4, 5]. Also, the symptoms become progressively worse, and much effort has been placed on the early diagnosis of the AD. Related to this, Mild Cognitive Impairment (MCI), defined as a transitional phase from cognitive changes of normal aging to those typically found in dementia, is an important construct [6]. Subjects with MCI present a high risk of developing AD, but still, most people with MCI will not progress to dementia (or AD) even after 10 years of follow-up [7, 8]. Thus, identifying MCI subjects who convert to AD can be crucial for the effective treatment of AD.

Neuroimaging techniques have shown promise as tools for presymptomatic AD detection [9, 10]. Much research has been focused on T1-weighted Magnetic Resonance Imaging (MRI). It is one of the most widely studied imaging techniques [11] because it is completely non-invasive, highly available, inexpensive compared to positron emission tomography and has an excellent contrast between different soft tissues. Over the past few years, many potential MRI markers, such as the whole brain, hippocampal, and entorhinal cortex atrophy, have been shown to have diagnostic value [12]. Also, these markers have been used as the features for Machine Learning (ML) algorithms trying to predict MCI-to-AD conversion.

Indeed, there has been a surge of proposed ML algorithms for automatically predicting the future conversion from MCI to AD based on MRI (e.g.,25 [13, 14, 15, 16]). This is partly driven by the free availability of large, high-quality datasets such as ADNI^1^. However, the principal focus has been in the development of new ML techniques, and their comparative evaluation has received much less attention. In particular, ML algorithms have used different types of feature sets extracted from MRI, including hippocampal volumes, volumes of the entorhinal cortex, cortical thickness measures, as well as voxel-based morphometry (VBM) features (e.g., [17, 18, 19, 20, 21] and [22] for a recent review). Despite that, systematic studies of advantages/disadvantages of various feature sets have been limited so far, and existing studies do not allow to make definite conclusions. To add to the confusion, high dimensional feature sets, such as cortical thickness or voxel-based morphometry, must be coupled with dimensionality reduction technique such as averaging the values within a brain region, Principal Component Analysis (PCA) or feature selection (see [23] for a review).

Existing comparisons between different feature representations do not provide a clear answer to the question we are interested in: “Is there a preferred representation of MRI for AD-conversion prediction?”. There are multiple reasons for this. The comparisons have been geared to theAD vs. control classification problem ([24, 25, 26]) they have not included voxel-based representations [24, 27], they have utilized very short follow-up (18-months [27, 28]), they have been based on a single learning algorithm [27, 29] and/or have had highly unbalanced pMCI and sMCI classes (in [29] 149 of 165 MCI subjects converted during the 4-year follow-up that is in stark contrast to conversion rates reported in other analyses [8]). An early and important study [28], which we want to highlight, compared various feature representations including hippocampal volumes, cortical thickness, and VBM with and without regional averaging. No feature representation in this study managed to perform significantly better than chance. This somewhat disappointing result could be because 1) the methods were early ones, mostly geared to the much easier normal control vs. AD sub ject classification problem, 2) the dataset was smaller than the one currently available, and 3) the MCI non-converter was somewhat arbitrarily defined as a subject who did not convert in 18 months period. Moradi et al. [30] evaluated their method over the same dataset as [28] managing to obtain significantly better performance than the chance level, pointing to the reason 1) as the most significant cause of the improvement.

Since [28], we can a find few studies of different feature representations presenting partially conflicting results. As an example, [21] found that the prognostic efficacy of hippocampus volumetry was better than combined regional volumetrics in two commercially available brain volumetric software packages for MCI conversion prediction. On the other hand, Gaser et al. [17] have demonstrated the superiority of their voxel-based brainAGE approach over the hippocampus volume biomarker and Westman et al. have emphasized the im-portance of having a complete set of regional features [27, 24]. Some researchers have opted to study feature selection, either supporting [20, 31] or opposingdata-driven feature selection. The comparisons of different automatic algorithms for hippocampal [33] and entorhinal cortex volumetry [34] have indicated that the algorithm-choice did not affect the classification accuracy. Intracranial volume adjustment to regional volumetry appears to only have subtle effects to the conversion prediction accuracy [35, 27]. Finally, it has been demonstrated that the neurophychological test scores are the best predictors of conversion, but combining them with MRI information leads to improved prediction accuracy [30, 36].

To close this information gap, we asked what type of feature representations are the best for the MRI-based AD-conversion prediction. We used follow-up period of 3 years to define the AD-conversion, twice longer than in [27, 28]. We evaluated the performance of various MRI features, including VBM-style, voxel-based features [17], coupled with feature preselection [30] or PCA-based dimensionality reduction, hippocampus volumes, volumes of the entorhinal cortex, and a complete set of regional volumetry, surface area, and cortical thickness measures extracted by FreeSurfer. This complements earlier studies which did not include voxels-based representations [27, 21]. We additionally evaluated age removal [30, 37], which have been found to improve the prognostic efficacy of ML-based MRI biomarkers. Moreover, we used two different classifiers (Support Vector Machines, SVM, and Regularized Logistic Regression, RLR) for reducing the classifier specificity of the conclusions and trained them applying a repeated 10-fold cross-Validation (CV) with a sound statistical inference to compare the methods, which can be seen as an improvement of separate training and test sets in [28].

## 2. Material and methods

### ADNI data

Data is collected from the the Alzheimers Disease Neuroimaging Initiative (ADNI) public database^2^. The ADNI initiative was launched in 2003 as a public-private partnership, led by Principal Investigator Michael W. Weiner, MD. The primary goal of ADNI has been to test whether serial magnetic resonance imaging (MRI), positron emission tomography (PET), other biological markers, and clinical and neuropsychological assessment can be combined to measure the progression of mild cognitive impairment (MCI) and early Alzheimers disease (AD)^3^.

ADNI material considered in this work included all subjects from ADNI1 for whom baseline MRI data (T1-weighted MP-RAGE sequence at 1.5 Tesla, typically 256 × 256 × 170 voxels with the voxel size of approximately 1 mm × 1 mm x× 1.2 mm) and sufficient follow-up information were available. We focused on the classification of MCI individuals based on their future diagnosis (AD or not AD) and, therefore, MRI scans were all obtained at the baseline visit.

Two flavors of this dataset were evaluated. The first dataset, Quality Control (QC) dataset, included 183 MCI subjects whose FreeSurfer 4.3 MRI segmentations had passed the complete quality control. The second one, Non QC dataset, included the complete dataset of 264 MCI subjects without any quality control. The reason for evaluating the two different sets was to study if the quality control yielded an improvement in the data analysis. Note that the QC dataset was a subset of the non-QC dataset.

Following [30], a subject was considered as a progressive MCI (pMCI) if diagnosed as MCI at baseline and the diagnosis changed to AD during the 3-year follow-up period. The subject was considered as a stable MCI (sMCI) if diagnosed as MCI at baseline and the diagnosis remained as MCI during the follow-up. The minimum length of follow-up was 3 years and the subject was excluded from the study if she converted after the 3 year follow-up, the diagnosis fluctuated after the 3-year follow-up period, or less than 3-years of follow-up information was available. Table 1 lists the main characteristics of the subjects of each dataset and the list of Roster IDs of the included subjects and their diagnostic categories are available in the supplement.

**Table 1:** Demographics of the two flavors of the dataset (QC and Non-QC) used in this work. The NC and AD subjects’ data were not used directly in the learning algorithms. The NC subjects were used for the age-correction. The AD and NC subjects were used in Moradi-method for feature selection.

|  | QC dataset |  |  |  | Non-QC dataset |  |  |  |
| --- | --- | --- | --- | --- | --- | --- | --- | --- |
|  | sMCI | pMCI | AD | NC | sMCI | pMCI | AD | NC |
| No. subjects | 73 | 110 | 126 | 182 | 100 | 164 | 200 | 231 |
| Males / Females | 46/27 | 58/52 | 65/61 | 91/91 | 66/34 | 97/67 | 103/97 | 119/112 |
| Age range | 59-88 | 55-89 | 55-91 | 60-90 | 57-88 | 55-89 | 55-91 | 60-90 |

### 2.1 Image preprocessing

Table 2 details the feature representations we investigated and their respective number of features. Hippocampus volumes consisted of left and right hippocampal volumes. Hippocampus + Entorhinal volumes consisted of left and 130 right volumes of hippocampus and left and right volumes of entorhinal cortex. We considered both raw volumes as well as volumes normalized by the intra-cranial volume (ICV) as it is still unclear if the normalization by ICV is beneficial for the prediction task [35, 27]. Region based features included a complete set of 257 regional cortical thickness, surface area and volume measures provided by FreeSurfer ^4, 5^,. We note that this set of features included also ICV.

**Table 2:**
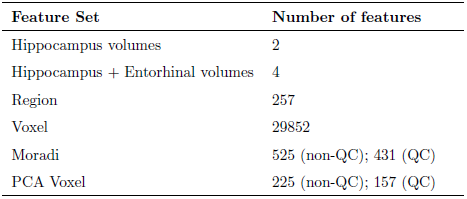
Summary of the sets of features considered in this study. Note that the number of Moradi and PCA voxel features was dataset dependent.

| Feature Set | Number of features |
| --- | --- |
| Hippocampus volumes | 2 |
| Hippocampus + Entorhinal volumes | 4 |
| Region | 257 |
| Voxel | 29852 |
| Moradi | 525 (non-QC); 431 (QC) |
| PCA Voxel | 225 (non-QC); 157 (QC) |

Freesurfer 4.3 software was employed for the extraction of hippocampus and entorhinal cortex volumes as well as region features. Particularly, the FreeSurfer 4.3 processing results available at the ADNI website were used (UCSF Cross-sectional Freesurfer version 4.3), and the description of the pipeline and the QC procedure can be found at^6^. The rationales for using the processing results provided by ADNI was to ensure that the processing pipeline was a standard one, the processing results are readily available to other researchers, and the quality control, independent from the authors of this study, has been performed. We note that albeit different versions of FreeSurfer can result in different segmentations, the classification results based on different software versions have been found to be the same [34].

Voxel-Based Morphometry (VBM) based features consisted of 29852 gray matter density values from the VBM style preprocessing by the VBM8 software. In brief, the MRIs were preprocessed into gray matter tissue images in the stereotactic space as described in [17, 30], smoothed with the 8-mm FWHM Gaussian kernel, resampled to 4 mm spatial resolution, and masked into 29852 voxels. In the Moradi set of features, VBM features were further processed through the feature selection method of [30]. This method applies MRIs of AD and NC subjects to select features for MCI classification through a repeated application of the elastic net penalized linear regression. We applied the ADNI data from 231 (182) normal controls and 200 (126) AD subjects for this feature selection with the non-QC (QC dataset). We reduced the number of VBM features also using principal component analysis (PCA). For this, we retained the PCA components that explained 90 % of the variance (see Tables 9, 10 and 11 of Supplementary Material for a performance comparison with different variance thresholds).

We further evaluated the representations with and without the age correction. The age correction may be important as the effects of normal aging on the brain structure partially overlap with the effects of AD [38, 37]. We applied the age correction procedure of [30]. This method estimates the age effect by a linear regression for each feature separately based on the MRIs of normal controls (231 normal controls with the age range from 55 to 90 years of ADNI) and then adjusts the features of the MCI subjects based on the estimated model.

### 2.2 Validation and test procedure

For the implementation and evaluation of the classification methods, we performed a repeated and nested 10-fold Cross Validation (CV). In the outer CV loop, data was split in 10 different folds from which one fold at time was designated as the test fold (for performance evaluation) and the nine remaining folds were used for classifier training. The train/test cycle was repeated with each fold once as the test fold. In the inner CV loop, each train fold was, itself, split into 10 validation folds from which one part was used to select the classifier hyperparameters. The optimal hyperparameters were selected evaluating either the classification accuracy (ACC, number of correctly classified samples over the total number of samples) or the Area Under the receiving operating Curve (AUC) [39]. It has been suggested that AUC has key advantages over ACC as a model selection criterion [40]. The nested CV was repeated 10 times, each with a different randomly selected folding scheme, to minimize the effect of a particular folding scheme to the results. Also, the hypothesis test we used to compare different representations requires the repeated use of CV.

To study the classifier performance, we considered several metrics: AUC, ACC, Sensitivity (SEN, number of correctly classified pMCI subjects divided by the total number of pMCI subjects) and Specificity (SPE, number of correctly classified sMCI subjects divided by the total number of sMCI subjects). We selected AUC as our principal performance measure as it is insensitive to the class-imbalance whereas ACC can be strongly affected by the class-imbalance.

### 2.3 Classifiers

To evaluate each feature set, we considered two types of widely used supervised learning classifiers: Support Vector Machine (SVM) [41] and elastic-net Regularized Logistic Regression (RLR) [42]. Accessible description of these learning methods can be found in [43]. For the SVM implementation, we used the Python open source library Scikit-learn^7^, which is based on the LIBSVM implementation^8^. For the RLR classifiers, we applied the GLMNET Python library^9^, which solves the resulting penalized optimization problem by a coordinate descent algorithm. We note that both of these learning algorithms tolerate high-dimensional data via regularization and are therefore suited for the cases where the number of features is higher than the number of subjects. Especially, elastic-net includes an L1-penalty, which leads to feature selection embedded to the classifier learning [44]. A large majority of supervised learning techniques have utilized these learning algorithms [22] and a comparison of different classification algorithms MCI-to-AD prediction is available in [45]. We note that we did not include Random Forests [46] as these are not straight-forwardly suitable for high-dimensional small-sample problems and the computation time and memory requirements for nearly all implementations would be prohibiting for the voxel-based features (however, see [47]).

For the case of the SVM classifier, we decided to use the linear SVM (we have also analyzed the possibility of using a RBF (Radial Basis Function) kernel, however, experimental results showed similar performances). In this way, we had to select only the soft margin parameter, *C*, whose value was explored among the set {10^−5^, 10^−4^, 10^−3^, 10^−2^, 10^−1^, 1, 10, 10^2^, 10^3^} (see [41] for notation). Despite considering the linear SVM, its implementation was carried out in the dual space, precomputing a linear kernel; in this way, we simplified the calculations and reduced the computation time with the high dimensional feature representations, such as VBM ones.

For the RLR classifier, using the notation of [42], we set the parameter of the elastic net *α* to 0.5, just in between lasso (*α*=1) and ridge (*α* = 0) regularization. The principal regularization parameter of the RLR (*λ*), which sets the balance between the regularization and the data terms, was chosen among the set of values {10^−10^, 10^−4^, 10^−3^, 5 · 10^−3^, 10^−2^, 5 · 10^−2^, 10^−1^, 5 · 10^−1^}. We previously demonstrated that selecting also the parameter *α* by cross-validation did not yield advantages over fixing its value to 0.5 [31]. However, we confirmed that this is the case with the setup of this paper by experimenting with different values of *α* (see Table 12 of the Supplementary Material).

Finally, as a step prior to training the classifiers, we normalized the data by removing its mean and scaling it to the unit variance.

### 2.4 Statistical test

To compare the AUC values provided by different approaches, we applied the corrected resampled t-test [48]. The problem in applying standard statistical methodology, such as uncorrected t-test to assess the differences between AUCs is that *r* × *k* AUC values from in a *k*-fold CV repeated *r* times are not statistically independent. Instead, the corrected resampled t-test assumes dependency among the AUCs in a *k*-fold CV repeated *r* times and, therefore, it allows to statistically compare two mean AUC values by correcting the variance estimation. The corrected resampled t-test can be seen as an improvement over the 5×2 CV of [49] and McNemar’s test for the classification accuracy [48]. Although the test was developed for the classification accuracy, it is as well applicable for testing the differences between AUCs.

To describe the test formally, let *n*_1_ and *n*_2_, respectively, denote the number of instances used for training and testing in each fold, and *a*_*ij*_ and *b*_*ij*_ represent the AUCs of the *i*-th fold and *j*-th run of the method *A* and *B* with *i* = 1, *…, k* and *j* = 1, *…, r*. Denoting the estimated mean and variance values of the differences between methods A and B by 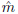 and 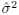 i.e.,

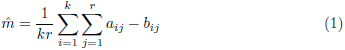

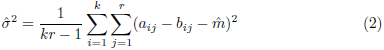

we can estimate the statistic of the test, *t*, as:

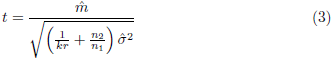

The statistic *t* follows a student’s t-distribution with *kr* − 1 degrees of freedom.

In our case, *r* = *k* = 10.

## 3. Results

Tables 3 – 8 show the results for the QC dataset and non-QC datasets using the AUC for model selection and the results of the non-QC dataset when the best model was selected using ACC. In particular, each table includes for the SVM and RLR classifiers the values of AUC, ACC, SEN, SPE, as well as three p-values from hypothesis tests comparing the AUCs: *p*_*Age*_ (comparing age removed features vs. non age removed features), *p*_*Hippo*_ (comparing hippocampus features with the remaining features for the age removed case) and *p*_*Class*_ (com-paring SVM vs. RLR results over the same set of features). The best results of each experimental setup, for each classifier and with/without the age correction process, have been marked in bold face. In addition, the standard deviation of each measure, computed as 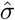 in Eq. (2), is included below its average value after *±* symbol.

**Table 3:** Cross-validated SVM performance measures with the QC dataset using AUC as the model selection criterion. Hippocampus (Hippo. and entor. vol.) volumes ICV refers to hippocampus (hippocampus + entorhinal) volumes normalized by the ICV.

| Feature Set | Age Removal | AUC (%) | ACC (%) | SEN (%) | SPE (%) | $p_{Age}$ | $p_{Hippo}$ | $p_{Class}$ |
| --- | --- | --- | --- | --- | --- | --- | --- | --- |
| Hippocampus | No | 73.50<br>$\pm 12.61$ | 63.15<br>$\pm 7.27$ | 93.00<br>$\pm 10.90$ | 18.16<br>$\pm 25.35$ | | | 0.873 |
| Hippocampus | Yes | 77.31<br>$\pm 11.22$ | 66.42<br>$\pm 9.67$ | 90.54<br>$\pm 11.78$ | 30.20<br>$\pm 31.18$ | 0.052 | | 0.819 |
| Hippocampus vol. ICV | No | 69.61<br>$\pm 12.60$ | 67.21<br>$\pm 9.88$ | 81.36<br>$\pm 9.91$ | 45.82<br>$\pm 14.42$ | | | 0.690 |
| Hippocampus vol. ICV | Yes | 72.67<br>$\pm 11.53$ | 68.31<br>$\pm 9.60$ | 82.36<br>$\pm 9.43$ | 47.11<br>$\pm 14.48$ | 0.124 | 0.169 | 0.416 |
| Hippo. and entor. | No | **75.91**<br>$\pm 11.84$ | 63.69<br>$\pm 7.91$ | 94.55<br>$\pm 9.09$ | 17.14<br>$\pm 25.78$ | | | 0.314 |
| Hippo. and entor. | Yes | **78.59%**<br>$\pm 10.84$ | 61.77<br>$\pm 5.64$ | 97.82<br>$\pm 7.02$ | 7.29<br>$\pm 21.19$ | 0.055 | 0.492 | 0.285 |
| Hippo. & entor. vol. ICV | No | 72.66<br>$\pm 11.26$ | 68.26<br>$\pm 10.27$ | 83.82<br>$\pm 10.17$ | 44.79<br>$\pm 16.58$ | | | 0.850 |
| Hippo. & entor. vol. ICV | Yes | 74.50<br>$\pm 10.65$ | 68.86<br>$\pm 9.84$ | 81.64<br>$\pm 9.70$ | 49.66<br>$\pm 15.61$ | 0.325 | 0.399 | 0.505 |
| Voxel features | No | 63.19<br>$\pm 12.21$ | 60.51<br>$\pm 5.66$ | 91.36<br>$\pm 14.74$ | 13.98<br>$\pm 21.14$ | | | 0.649 |
| Voxel features | Yes | 66.67<br>$\pm 11.71$ | 61.65<br>$\pm 5.69$ | 91.82<br>$\pm 12.23$ | 16.20<br>$\pm 17.19$ | 0.175 | 0.042 | 0.923 |
| PCA VF 90 | No | 64.33<br>$\pm 11.63$ | 60.11<br>$\pm 5.16$ | 90.36<br>$\pm 14.48$ | 14.38<br>$\pm 20.84$ | | | 0.791 |
| PCA VF 90 | Yes | 66.78<br>$\pm 11.67$ | 61.09<br>$\pm 5.74$ | 91.82<br>$\pm 14.35$ | 14.64<br>$\pm 22.43$ | 0.351 | 0.042 | 0.569 |
| Moradi features | No | 71.92<br>$\pm 10.62$ | 63.54<br>$\pm 7.69$ | 92.63<br>$\pm 10.91$ | 19.66<br>$\pm 23.45$ | | | 0.025 |
| Moradi features | Yes | 75.08<br>$\pm 9.80$ | 62.32<br>$\pm 5.60$ | 97.10<br>$\pm 7.03$ | 9.86<br>$\pm 20.64$ | 0.243 | 0.610 | 0.001 |
| Region features | No | 74.06<br>$\pm 12.21$ | 64.67<br>$\pm 7.29$ | 92.27<br>$\pm 11.31$ | 22.91<br>$\pm 24.08$ | | | 0.739 |
| Region features | Yes | 77.34<br>$\pm 10.22$ | 69.29<br>$\pm 8.96$ | 88.45<br>$\pm 10.92$ | 37.27<br>$\pm 26.62$ | 0.241 | 0.990 | 0.759 |

**Table 4:** Cross-validated RLR performance measures with the QC dataset using AUC as the model selection criterion. Hippocampus (Hippo. and entor. vol.) volumes ICV refers to hippocampus (hippocampus + entorhinal) volumes normalized by the ICV.

| Feature Set | Age Removal | AUC (%) | ACC(%) | SEN(%) | SPE(%) | $p_{Age}$ | $p_{Hippo}$ | $p_{Class}$ |
| --- | --- | --- | --- | --- | --- | --- | --- | --- |
| Hippocampus | No | 73.38<br>$\pm 12.72$ | 68.41<br>$\pm 10.22$ | 82.00<br>$\pm 11.85$ | 47.86<br>$\pm 17.66$ | | | |
| Hippocampus | Yes | 77.19<br>$\pm 11.15$ | 71.31<br>$\pm 9.16$ | 83.91<br>$\pm 10.82$ | 52.43<br>$\pm 16.82$ | 0.042 | | |
| Hippocampus vol. ICV | No | 69.80<br>$\pm 12.67$ | 66.57<br>$\pm 9.47$ | 75.16<br>$\pm 11.87$ | 38.88<br>$\pm 18.27$ | | | |
| Hippocampus vol. ICV | Yes | 72.25<br>$\pm 11.66$ | 68.32<br>$\pm 9.29$ | 84.64<br>$\pm 11.26$ | 43.68<br>$\pm 16.23$ | 0.253 | 0.138 | |
| Hippo. and entor. | No | 72.52<br>$\pm 12.18$ | 66.85<br>$\pm 9.54$ | 80.82<br>$\pm 10.33$ | 45.73<br>$\pm 18.48$ | | | |
| Hippo. and entor. | Yes | 74.13<br>$\pm 11.28 \%$ | 68.05<br>$\pm 9.62$ | 81.64<br>$\pm 10.40$ | 47.55<br>$\pm 18.97$ | 0.085 | 0.767 | |
| Hippo. & entor. vol. ICV | No | 71.72<br>$\pm 11.38$ | 68.59<br>$\pm 9.89 \%$ | 84.47<br>$\pm 11.85$ | 42.60<br>$\pm 17.11$ | | | |
| Hippo. & entor. vol. ICV | Yes | 73.60<br>$\pm 10.83 \%$ | 68.90<br>$\pm 9.23$ | 83.51<br>$\pm 11.50$ | 45.00<br>$\pm 18.30$ | 0.332 | 0.348 | |
| Voxel features | No | 64.40<br>$\pm 11.95 \%$ | 61.91<br>$\pm 9.86$ | 76.18<br>$\pm 13.22$ | 40.43<br>$\pm 15.45$ | | | |
| Voxel features | Yes | 66.94<br>$\pm 11.15$ | 63.57<br>$\pm 9.48$ | 77.36<br>$\pm 13.20$ | 40.43<br>$\pm 16.37$ | 0.268 | 0.029 | |
| PCA VF 90 | No | 63.63<br>$\pm 11.20$ | 60.88<br>$\pm 8.33$ | 79.45<br>$\pm 14.00$ | 33.00<br>$\pm 18.91$ | | | |
| PCA VF 90 | Yes | 65.35<br>$\pm 12.33$ | 61.28<br>$\pm 8.44$ | 81.73<br>$\pm 14.46$ | 30.43<br>$\pm 19.14$ | 0.573 | 0.018 | |
| Moradi features | No | 65.83<br>$\pm 11.36$ | 63.28<br>$\pm 9.68$ | 75.00<br>$\pm 13.01$ | 42.73<br>$\pm 16.84$ | | | |
| Moradi features | Yes | 67.89<br>$\pm 10.43$ | 65.52<br>$\pm 9.71$ | 77.54<br>$\pm 13.69$ | 57.50<br>$\pm 17.24$ | 0.348 | 0.038 | |
| Region features | No | **74.57**<br>$\pm 10.92$ | 70.67<br>$\pm 8.92$ | 81.91<br>$\pm 10.40$ | 53.70<br>$\pm 17.31$ | | | |
| Region features | Yes | **77.91**<br>$\pm 9.59$ | 71.93<br>$\pm 9.22$ | 81.45<br>$\pm 10.90\%$ | 57.50<br>$\pm 18.26$ | 0.171 | 0.839 | |

The AUC values of region features were the highest in all the experiments. However, the performance improvement over the hippocampus feature set, which was our baseline, did not reach the statistical significance and these improved AUCs need to be interpreted with care. In the particular case of the non-QC dataset and the RLR classifier, the regions feature set produced significantly higher AUC than hippocampus volumes.

Figure 1 depicts the ROC curves for the different feature sets under study for the RLR classifier in the non-QC dataset. Focusing on the center of these curves (see the panel 1 b), we can corroborate that the region feature set appeared superior, but the performance differences were small. To avoid crowding, the ROCs of the PCA voxel feature set were not visualized as PCA voxel features always performed worse than the voxel features without PCA. For similar reason, the figure only displays the ROC curves of raw Hippocampus and Hippocampus + Entorhinal cortex volumes and not the ICV-normalized ones. The same principle will be followed in later figures.

**Figure 1:**
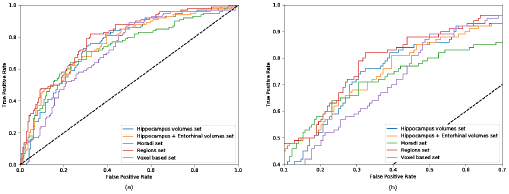
ROC curves corresponding to distinct the features sets used in RLR classification with the non-QC dataset. The age effect was removed.

Regarding the use of two different classifiers, differences between AUCs of SVM and RLR were not significant. However, SVM yielded low specificity values and the relation between SPE and SEN was more balanced with the RLR classifier. Because of this we studied whether the use of AUC as the model selection criteria contributed to this imbalance with the SVM classifier. Using ACC as a model selection criterion notably reduced this SPE/SEN imbalance as can be seen in Figure 2 where the specificity values are compared between ACC and AUC based model selection. As the comparison of Tables 5 - 8 reveals, the final AUC values did not markedly differ between the two model selectors. With ACC as the model selection criterion, the sensitivity values were still markedly higher than the specificity values. Some insight to the phenomenon can be obtained by visual analysis of the Hippocampus feature set, with just 2 features, thereby permitting visual analysis. Figure 3 shows the datapoints along with the decision regions for two classes over 10 CV folds. It can be observed that the data from two classes were highly overlapping, and in these cases, the classification boundary has a tendency to shift more towards the majority class (pMCI in this case) than what might be expected based on a modest class-imbalance.

**Table 5:**
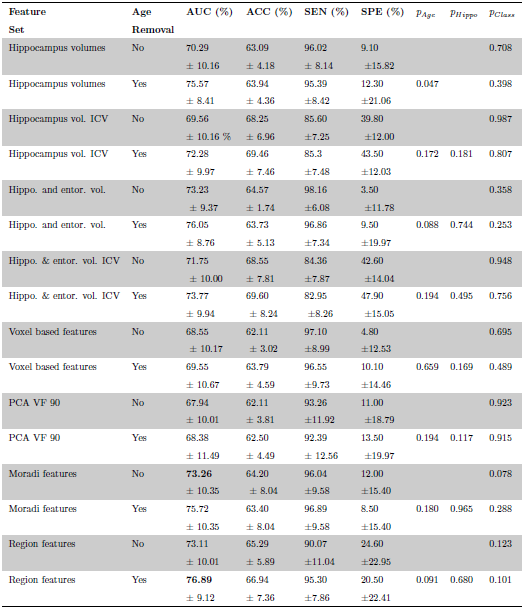
Cross-validated SVM performance measures with the non QC dataset using AUC as the model selection criterion. Hippocampus (Hippo. and entor. vol.) volumes ICV refers to hippocampus (hippocampus + entorhinal) volumes normalized by the ICV.

**Table 6:** Cross-validated RLR performance measures with the non QC dataset using AUC as the model selection criterion. Hippocampus (Hippo. and entor. vol.) volumes ICV refers to hippocampus (hippocampus + entorhinal) volumes normalized by the ICV.

| Feature Set | Age Removal | AUC (%) | ACC (%) | SEN (%) | SPE (%) | $p_{age}$ | $p_{Hippo}$ | $p_{Class}$ |
| --- | --- | --- | --- | --- | --- | --- | --- | --- |
| Hippocampus volumes | No | 70.95<br>$\pm 8.89$ | 67.00<br>$\pm 7.29$ | 88.04<br>$\pm 10.43$ | 32.40<br>$\pm 15.24$ | | | |
| Hippocampus volumes | Yes | 74.95<br>$\pm 8.25$ | 67.68<br>$\pm 7.00$ | 86.34<br>$\pm 8.98$ | 37.00<br>$\pm 13.60$ | 0.046 | | |
| Hippocampus vol. ICV | No | 69.60<br>$\pm 10.41$ | 67.94<br>$\pm 6.88$ | 87.91<br>$\pm 8.49\%$ | 35.20<br>$\pm 15.84$ | | | |
| Hippocampus vol. ICV | Yes | 72.37<br>$\pm 10.03$ | 69.44<br>$\pm 7.47$ | 88.76<br>$\pm 8.12$ | 37.80<br>$\pm 15.20$ | 0.148 | 0.272 | |
| Hippo. and entor. vol. | No | 72.59<br>$\pm 9.23$ | 69.72<br>$\pm 7.86$ | 85.88<br>$\pm 9.59\%$ | 43.20<br>$\pm 15.35$ | | | |
| Hippo. and entor. vol. | Yes | 75.31<br>$\pm 9.03$ | 70.61<br>$\pm 7.47$ | 84.98<br>$\pm 9.94$ | 47.00<br>$\pm 14.93$ | 0.094 | 0.801 | |
| Hippo. & entor. vol. ICV | No | 71.72<br>$\pm 9.76$ | 68.59<br>$\pm 8.08$ | 84.47<br>$\pm 9.24$ | 42.60<br>$\pm 16.71$ | | | |
| Hippo. & entor. vol. ICV | Yes | 73.60<br>$\pm 9.93$ | 68.90<br>$\pm 7.63$ | 83.51<br>$\pm 8.88$ | 45.00<br>$\pm 16.03$ | 0.210 | 0.603 | |
| Voxel based features | No | 69.46<br>$\pm 9.69$ | 66.63<br>$\pm 7.79$ | 83.42<br>$\pm 8.39$ | 39.10<br>$\pm 15.88$ | | | |
| Voxel based features | Yes | 71.34<br>$\pm 10.34$ | 66.99<br>$\pm 8.04$ | 82.68<br>$\pm 9.58$ | 41.30<br>$\pm 15.40$ | 0.370 | 0.394 | |
| PCA VF 90 | No | 67.75<br>$\pm 10.10$ | 65.19<br>$\pm 8.09$ | 84.09<br>$\pm 11.33$ | 34.20<br>$\pm 17.33$ | | | |
| PCA VF 90 | Yes | 68.63<br>$\pm 10.37$ | 65.25<br>$\pm 7.02$ | 85.72<br>$\pm 10.96$ | 31.70<br>$\pm 15.81$ | 0.713 | 0.116 | |
| Moradi features | No | 69.94<br>$\pm 10.05$ | 67.77<br>$\pm 7.77$ | 83.81<br>$\pm 10.11$ | 41.50<br>$\pm 14.79$ | | | |
| Moradi features | Yes | 74.04<br>$\pm 9.37$ | 70.84<br>$\pm 7.28$ | 86.79<br>$\pm 8.26$ | 44.70<br>$\pm 14.66$ | 0.068 | 0.798 | |
| Region features | No | **76.38**<br>$\pm 8.67$ | 71.27<br>$\pm 7.53$ | 85.97<br>$\pm 9.57$ | 47.10<br>$\pm 16.33$ | | | |
| Region features | Yes | **79.58**<br>$\pm 7.71$ | 71.73<br>$\pm 7.56$ | 84.07<br>$\pm 9.26$ | 51.50<br>$\pm 15.58$ | 0.120 | 0.060 | |

**Figure 2:**
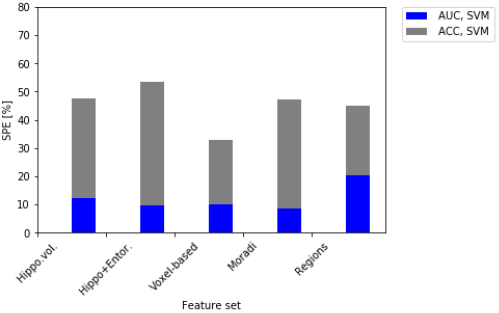
Specificity values of SVM classifiers when AUC and ACC were used for model selection. The models selected with ACC resulted in specificity values close 50 % whereas the models selected with AUC resulted in very low specificity values.

**Figure 3:**
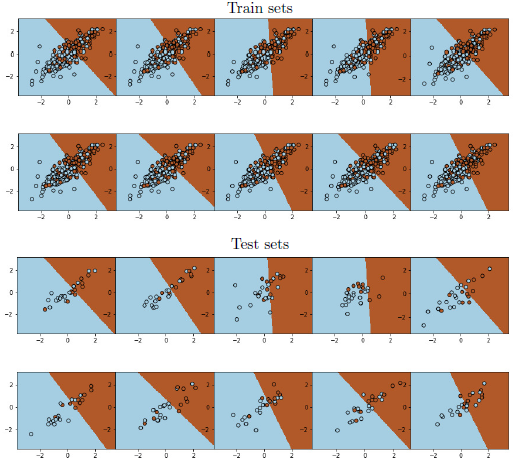
SVM classification boundaries with the age corrected hippocampus non-QC feature set overlaid to the train and test data. ACC was used as the model selection criteria. Each panel depicts the classifier training in a single CV fold (folds from the first CV run are shown). On top of the decision regions, train or test sets of that particular fold are plotted. Red color corresponds to sMCI class and blue corresponds to pMCI class. Note that the decision regions are always based on the training set, and therefore they are the same whether overlaying test or train data. x-axis (y-axis) of the feature space corresponds to right (left) Hippocampus volume. The feature values are normalized as explained in Section 2.3.

We evaluated the effects of age removal on the feature sets. For this purpose, Figure 4 shows a detailed analysis of the advantages of removing the age effects. As a result, classification scores improved for every age removed effects feature set (see the panel 4 c). However, as visible in Tables 3-6, significant improve-ment (p-value < 0.1) was observed only for hippocampus and hippocampus + entorhinal volume feature sets.

**Figure 4:**
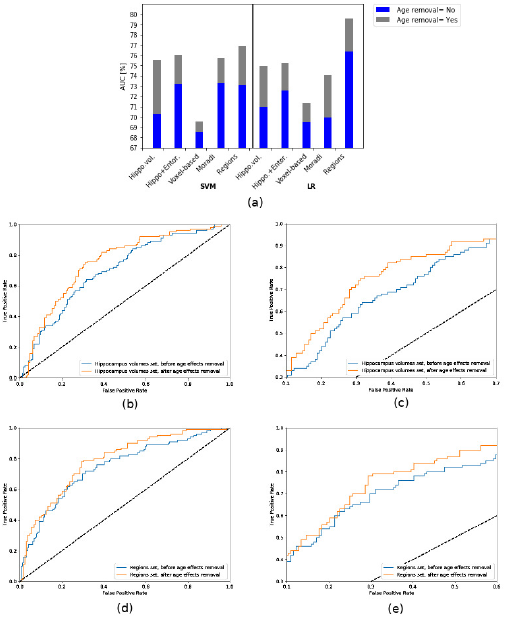
Analysis of age removal effects: (a) AUC comparison for different feature sets and both classifiers; (b) and (c) ROC curves for RLR classifier using hippocampus volumes; (d) and (e) ROC curves for RLR classifier using region features. Age removal improved predictions in all cases.

The differences between the AUCs of raw and ICV-normalized hippocampus and hippocampus + entorhinal volumes were not significant. Surprisingly, the raw volumes performed slightly better in terms of AUC within each dataset. However, this result agrees with findings in [35, 27] and it is not central for the purposes of this work to analyze the potential reasons for this result.

Finally, Figure 5 shows the differences between QC and non QC datasets when age effects were removed. As expected Hippocampus and Hippocampus plus Entorhinal volumes were benefited from the Quality Control process, whereas remaining features sets resulted in better performances when all the available data were used.

**Figure 5:**
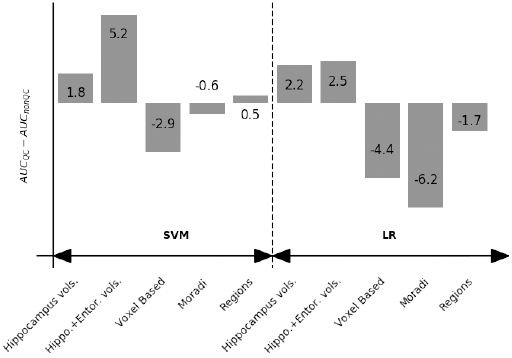
Differences between the AUC values with the QC dataset and the non-QC dataset for SVM (left) and RLR (right).

## 4. Discussion

In this work, we compared six different feature representations of MRI for predicting the AD conversion in MCI subjects. The feature sets we studied varied from high dimensional feature sets produced by VBM via regional cortical thickness, surface area, and volumetry to simple and easily interpretable features such as hippocampus and entorhinal cortex volumes (see Table 2). We addressed the feature representations using two learning algorithms, SVM and RLR, and with several metrics, AUC, ACC, SEN and SPE, that gave a reliable insight into the relative performance of different feature sets. AUC was selected as the principal figure of merit, due to its insensitivity to the class imbalance (note that the datasets contained twice the number of pMCIs (subjects who converted to AD) compared to sMCIs (subjects who remained as MCIs)). The evaluation process was carried out with a nested 10-fold CV repeated 10 times ensuring the insensitivity of the conclusions to random train/test division of the holdout method used previously [28]. Selecting the parameters of the classifiers inside nested CV ensures that there are no biases towards particular feature representations due to arbitrarily selected classifier parameters

We found that age-corrected *regions feature set*^10^ outperformed the remaining feature sets, specifically in AUC, even though the improvement did not reach statistical significance. This result suggests that regions based features were equal or better predictors than the left and right hippocampal volumes (HV) alone (which were included in the region feature set). This is interesting as a recent study [21] concluded that HV had the highest AUC among a set of individual regional volume features and was better in terms of the prognostic efficacy of combining various volumetrics. Their experimental setting was similar to the one analyzed here, however, with three main differences. First, removing age related effects from MRI data was not considered; second, the set of pMCI patients was about half of ours; and, third, the combined volumetric analysis did not consider measures such as surface area or cortical thickness. This can explain the improvement in the best classification accuracy from 69 % of [21] to 80 % in the present study.

Voxel-based representations did not perform well in this study when coupled with standard feature reduction techniques (elastic-net or PCA). This was in contrast to a recent data-analysis competition, where the goal was to classify subjects into NC, MCI, and AD categories based on MRI [26]. However, as multiple factors have effect to a performance of an approach in a data analysis competition, definite conclusions on feature representations cannot be made based on such competitions. However, also in our own experience, voxel-based methods, coupled with elastic-net feature selection, perform well in classifying between NC and AD or NC and MCI [31]. These discrepancies may suggest that NC vs. MCI (or AD) classification and AD-conversion prediction have different characteristics. Further, we note that feature pre-selection based on AD and NC data suggested by Moradi et al. [30] improved the conversion prediction accuracy markedly.

Retico et al. found that the voxel based VBM features best discriminate between sMCI and pMCI after applying Recursive Feature Elimination (RFE) [20]. However, again, the maximum accuracy in [20] was much lower than the accuracies in the present study and pMCI vs. sMCI classifiers were trained only using AD and NC subjects that may explain this. Additionally, the statistical framework was incomplete as no hypothesis testing was done and the exact definition of stable MCI class remained unclear. Other works, such as [18], concluded that the combination of different feature representations resulted into a better classification accuracy than one representation alone. Again, the classification accuracies were lower than in the present work. Moreover, [18] selected classifier hyperparameters based on test data that may cause upward bias in the reported accuracies [15].

It is important to point out that while our classification accuracies were better than those in the studies reviewed above, the performance measures are not directly comparable because different definitions of pMCI and sMCI. In fact, this is a problem that complicates the comparison of ML methods for this particular application and it is reviewed in further length in [22]. Namely, the definition of sMCI subject based on a certain cutoff (say 3 years) is problematic as this simple criterion would place a subject who received an AD diagnosis 4 years after the baseline visit into the sMCI category. Our view is that this would create unrealistic heterogeneity into the sMCI class and therefore tracking subjects’ status after the cutoff is necessary (if possible). We have populated our sMCI category based on all the information available by ADNI.

Regarding the used ML methods, RLR provided, in general, similar AUC values than SVM, but had an advantage of higher specificity (it classified sMCI cases much better than the SVM did). SVM had a tendency of overpopulating the pMCI class. However, in the case of SVM, low specificity seemed to depend on the using AUC as the criterion for the hyperparameter selection. The values in Tables 7 and 8 reveal how selecting the hyperparameters instead through ACC resulted in an overall improvement of specificity with a small loss of sensitivity. This is an interesting phenomenon, as it seems to be a problem of a specific class of learning algorithms, which invites further research. However, as this issue is not central to the goals of this work, we do not analyze it further. Also with the ACC model selection and with RLR, the specificity values were lower than the sensitivity values. However, as already mentioned (see Fig. 3), this level of SEN/SPE imbalance can be explained by the slight class imbalance (approximately 60 % pMCI and 40 % sMCI) and overlapping feature densities.

**Table 7:**
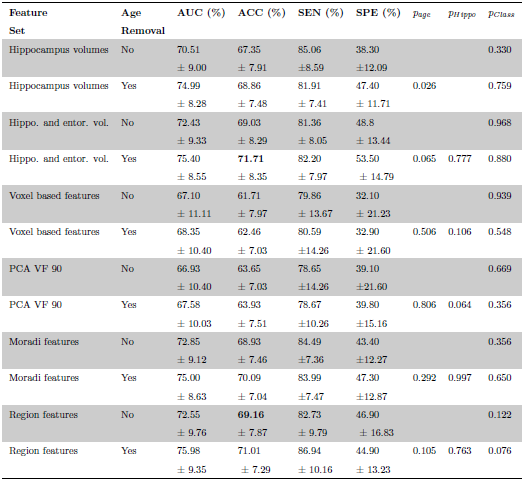
Cross-validated SVM performance measures with the non-QC dataset using ACC as the model selection criterion.

| Feature Set | Age Removal | AUC (%) | ACC (%) | SEN (%) | SPE (%) | $p_{age}$ | $p_{Hippo}$ | $p_{Class}$ |
| --- | --- | --- | --- | --- | --- | --- | --- | --- |
| Hippocampus volumes | No | 70.51<br>$\pm 9.00$ | 67.35<br>$\pm 7.91$ | 85.06<br>$\pm 8.59$ | 38.30<br>$\pm 12.09$ | | | 0.330 |
| Hippocampus volumes | Yes | 74.99<br>$\pm 8.28$ | 68.86<br>$\pm 7.48$ | 81.91<br>$\pm 7.41$ | 47.40<br>$\pm 11.71$ | 0.026 | | 0.759 |
| Hippo. and entor. vol. | No | 72.43<br>$\pm 9.33$ | 69.03<br>$\pm 8.29$ | 81.36<br>$\pm 8.05$ | 48.8<br>$\pm 13.44$ | | | 0.968 |
| Hippo. and entor. vol. | Yes | 75.40<br>$\pm 8.55$ | **71.71**<br>$\pm 8.35$ | 82.20<br>$\pm 7.97$ | 53.50<br>$\pm 14.79$ | 0.065 | 0.777 | 0.880 |
| Voxel based features | No | 67.10<br>$\pm 11.11$ | 61.71<br>$\pm 7.97$ | 79.86<br>$\pm 13.67$ | 32.10<br>$\pm 21.23$ | | | 0.939 |
| Voxel based features | Yes | 68.35<br>$\pm 10.40$ | 62.46<br>$\pm 7.03$ | 80.59<br>$\pm 14.26$ | 32.90<br>$\pm 21.60$ | 0.506 | 0.106 | 0.548 |
| PCA VF 90 | No | 66.93<br>$\pm 10.40$ | 63.65<br>$\pm 7.03$ | 78.65<br>$\pm 14.26$ | 39.10<br>$\pm 21.60$ | | | 0.669 |
| PCA VF 90 | Yes | 67.58<br>$\pm 10.03$ | 63.93<br>$\pm 7.51$ | 78.67<br>$\pm 10.26$ | 39.80<br>$\pm 15.16$ | 0.806 | 0.064 | 0.356 |
| Moradi features | No | 72.85<br>$\pm 9.12$ | 68.93<br>$\pm 7.46$ | 84.49<br>$\pm 7.36$ | 43.40<br>$\pm 12.27$ | | | 0.356 |
| Moradi features | Yes | 75.00<br>$\pm 8.63$ | 70.09<br>$\pm 7.04$ | 83.99<br>$\pm 7.47$ | 47.30<br>$\pm 12.87$ | 0.292 | 0.997 | 0.650 |
| Region features | No | 72.55<br>$\pm 9.76$ | **69.16**<br>$\pm 7.87$ | 82.73<br>$\pm 9.79$ | 46.90<br>$\pm 16.83$ | | | 0.122 |
| Region features | Yes | 75.98<br>$\pm 9.35$ | 71.01<br>$\pm 7.29$ | 86.94<br>$\pm 10.16$ | 44.90<br>$\pm 13.23$ | 0.105 | 0.763 | 0.076 |

**Table 8:** Cross-validated RLR performance measures with the non-QC dataset using ACC as the model selection criterion.

| Feature Set | Age Removal | AUC (%) | ACC (%) | SEN (%) | SPE (%) | $p_{age}$ | $p_{Hippo}$ | $p_{Class}$ |
| --- | --- | --- | --- | --- | --- | --- | --- | --- |
| Hippocampus volumes | No | 70.96<br>$\pm 9.07$ | 66.28<br>$\pm 7.21$ | 88.92<br>$\pm 10.73$ | 29.10<br>$\pm 13.94$ | | | |
| Hippocampus volumes | Yes | 74.85<br>$\pm 8.39$ | 69.12<br>$\pm 7.31$ | 84.10<br>$\pm 9.25$ | 44.50<br>$\pm 14.24$ | 0.050 | | |
| Hippo. and entor. vol. | No | 72.40<br>$\pm 9.15$ | 69.55<br>$\pm 7.90$ | 85.50<br>$\pm 9.18$ | 43.30<br>$\pm 15.81$ | | | |
| Hippo. and entor. vol. | Yes | 75.50<br>$\pm 9.11$ | 70.50<br>$\pm 7.54$ | 84.68<br>$\pm 9.50$ | 47.20<br>$\pm 15.69$ | 0.056 | 0.650 | |
| Voxel based features | No | 67.33<br>$\pm 10.03$ | 64.81<br>$\pm 8.03$ | 80.76<br>$\pm 9.45$ | 38.60<br>$\pm 16.43$ | | | |
| Voxel based features | Yes | 69.95<br>$\pm 10.43$ | 66.15<br>$\pm 8.26$ | 80.21<br>$\pm 10.41$ | 43.10<br>$\pm 16.35$ | 0.235 | 0.244 | |
| PCA VF 90 | No | 68.01<br>$\pm 10.43$ | 64.95<br>$\pm 8.26$ | 86.20<br>$\pm 10.41$ | 30.10<br>$\pm 16.35$ | | | |
| PCA VF 90 | Yes | 69.63<br>$\pm 9.55$ | 65.08<br>$\pm 7.75$ | 84.72<br>$\pm 13.08$ | 32.90<br>$\pm 17.68$ | 0.484 | 0.180 | |
| Moradi features | No | 71.08<br>$\pm 9.55$ | 68.60<br>$\pm 6.59$ | 85.35<br>$\pm 8.80$ | 41.10<br>$\pm 14.83$ | | | |
| Moradi features | Yes | 74.10<br>$\pm 9.33$ | 70.42<br>$\pm 7.38$ | 85.94<br>$\pm 9.51$ | 45.00<br>$\pm 14.39$ | 0.125 | 0.836 | |
| Region features | No | 75.91<br>$\pm 9.12$ | **71.01**<br>$\pm 7.89$ | 84.52<br>$\pm 9.27$ | 48.80<br>$\pm 16.02$ | | | |
| Region features | Yes | 79.41<br>$\pm 7.98$ | **72.07**<br>$\pm 7.81$ | 84.24<br>$\pm 9.40$ | 52.10<br>$\pm 15.32$ | 0.123 | 0.717 | |

p-values from hypothesis tests comparing the AUCs: $p_{Age}$ (comparing age removed features vs. non age removed features), $p_{Hippo}$ (comparing hippocampus features with the remaining features for the age removed case) and $p_{Class}$ (comparing SVM vs. RLR results over the same set of features). The best results of each experimental setup, for each classifier and with/without the age correction process, have been marked in bold face. In addition, the standard deviation of each measure, computed as $\hat{\sigma}$ in Eq. (2), is included below its average value after ± symbol.

There were no significant differences between the classification accuracies or AUCs obtained with non-QC and QC datasets. However, the small differences between the two datasets were as expected as shown in Figure 5. For Hippocampus and Hippocampus and Entorhinal volumes, the QC was moderately useful whereas for the Moradi and Voxel based features it was moderately detrimental. This is as expected since the QC was based on Freesurfer segmentations (as Hippocampus and Entorhinal volumes) but the voxel-based and Moradi features were not. Interestingly, for region based features (also based on Freesurfer segmentation), the QC seemed not to influence the performance of the classifier.

It is remarkable that the age removal seem to be a key for better performances. As Figure 4 illustrates, age removal always led to better classification performances, although the improvements were not always statistically significant. This agrees with a recent work of [31] which demonstrated the same for NC vs. MCI classification.

## 5. Conclusion

This paper evaluated the performance of various types of MRI features for the future AD conversion prediction and it also analyzed the performance of each feature set over two classifiers (Support Vector Machines and Regularized Logistic Regression) and with and without applying an age correction process.

Experimental results showed that regional features consistently yielded the best performance, although the performance difference to other features was not statistically significant. Besides, the age removal seemed to be a key for better performances, but the improvement reached statistical significance only rarely.

## 6. Supplementary Material

**Table 9:** Cross-validated performance measures of the PCA voxel feature set with the QC dataset using AUC as the model selection criterion.

| Classifier | Feature Set | Age Removal | AUC | ACC | SEN | SPE | $p_{Age}$ | $p_{Hippo}$ | $p_{Class}$ |
| --- | --- | --- | --- | --- | --- | --- | --- | --- | --- |
| SVM | PCA VF 99 | No | 63.50 % | 59.88 % | 92.73 % | 10.23 % |  |  | 0.604 |
| SVM | PCA VF 99 | Yes | 65.53 % | 60.75 % | 92.64 % | 12.54 % | 0.535 | 0.035 | 0.509 |
| SVM | PCA VF 95 | No | 64.22 % | 60.75 % | 94.27 % | 10.11 % |  |  | 0.826 |
| SVM | PCA VF 95 | Yes | 66.81 % | 60.74 % | 93.27 % | 11.59 % | 0.332 | 0.045 | 0.556 |
| SVM | PCA VF 90 | No | 64.33 % | 60.11 % | 90.36 % | 14.38 % |  |  | 0.791 |
| SVM | PCA VF 90 | Yes | 66.78 % | 61.09 % | 91.82 % | 14.64 % | 0.351 | 0.042 | 0.569 |
| SVM | PCA VF 75 | No | 63.08 % | 59.81 % | 84.45 % | 22.57 % |  |  | 0.773 |
| SVM | PCA VF 75 | Yes | 66.89 % | 62.25 % | 87.45 % | 24.45 % | 0.229 | 0.062 | 0.395 |
| LR | PCA VF 99 | No | 61.95 % | 59.12 % | 78.73 % | 26.68 % |  |  |  |
| LR | PCA VF 99 | Yes | 63.40 % | 61.33 % | 80.82 % | 31.91 % | 0.600 | 0.006 |  |
| LR | PCA VF 95 | No | 63.59 % | 61.19 % | 80.00 % | 32.93 % |  |  |  |
| LR | PCA VF 95 | Yes | 65.20 % | 61.22 % | 80.64 % | 31.91 % | 0.549 | 0.011 |  |
| LR | PCA VF 90 | No | 63.63 % | 60.88 % | 79.45 % | 33.00 % |  |  |  |
| LR | PCA VF 90 | Yes | 65.35 % | 61.28 % | 81.73 % | 30.43 % | 0.573 | 0.018 |  |
| LR | PCA VF 75 | No | 63.83 % | 61.08 % | 74.91 % | 40.30 % |  |  |  |
| LR | PCA VF 75 | Yes | 64.90 % | 61.69 % | 76.27 % | 39.68 % | 0.722 | 0.013 |  |

**Table 10:**
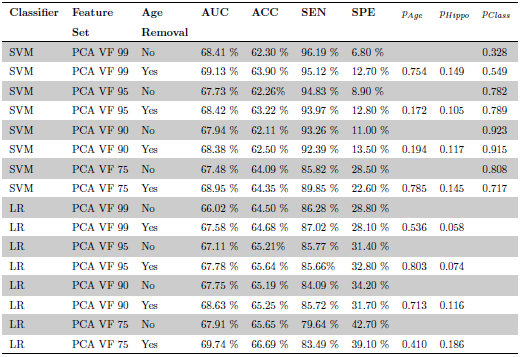
Cross-validated performance measures of the PCA voxel feature set with the non QC dataset using AUC as the model selection criterion

| Classifier | Feature Set | Age Removal | AUC | ACC | SEN | SPE | $p_{Age}$ | $p_{Hippo}$ | $p_{Class}$ |
| --- | --- | --- | --- | --- | --- | --- | --- | --- | --- |
| SVM | PCA VF 99 | No | 68.41 % | 62.30 % | 96.19 % | 6.80 % |  |  | 0.328 |
| SVM | PCA VF 99 | Yes | 69.13 % | 63.90 % | 95.12 % | 12.70 % | 0.754 | 0.149 | 0.549 |
| SVM | PCA VF 95 | No | 67.73 % | 62.26% | 94.83 % | 8.90 % |  |  | 0.782 |
| SVM | PCA VF 95 | Yes | 68.42 % | 63.22 % | 93.97 % | 12.80 % | 0.172 | 0.105 | 0.789 |
| SVM | PCA VF 90 | No | 67.94 % | 62.11 % | 93.26 % | 11.00 % |  |  | 0.923 |
| SVM | PCA VF 90 | Yes | 68.38 % | 62.50 % | 92.39 % | 13.50 % | 0.194 | 0.117 | 0.915 |
| SVM | PCA VF 75 | No | 67.48 % | 64.09 % | 85.82 % | 28.50 % |  |  | 0.808 |
| SVM | PCA VF 75 | Yes | 68.95 % | 64.35 % | 89.85 % | 22.60 % | 0.785 | 0.145 | 0.717 |
| LR | PCA VF 99 | No | 66.02 % | 64.50 % | 86.28 % | 28.80 % |  |  |  |
| LR | PCA VF 99 | Yes | 67.58 % | 64.68 % | 87.02 % | 28.10 % | 0.536 | 0.058 |  |
| LR | PCA VF 95 | No | 67.11 % | 65.21% | 85.77 % | 31.40 % |  |  |  |
| LR | PCA VF 95 | Yes | 67.78 % | 65.64 % | 85.66% | 32.80 % | 0.803 | 0.074 |  |
| LR | PCA VF 90 | No | 67.75 % | 65.19 % | 84.09 % | 34.20 % |  |  |  |
| LR | PCA VF 90 | Yes | 68.63 % | 65.25 % | 85.72 % | 31.70 % | 0.713 | 0.116 |  |
| LR | PCA VF 75 | No | 67.91 % | 65.65 % | 79.64 % | 42.70 % |  |  |  |
| LR | PCA VF 75 | Yes | 69.74 % | 66.69 % | 83.49 % | 39.10 % | 0.410 | 0.186 |  |

**Table 11:** Cross-validated performance measures of the PCA voxel feature set with the non-QC dataset using ACC as the model selection criterion

| Classifier | Feature Set | Age Removal | AUC | ACC | SEN | SPE | $p_{Age}$ | $p_{Hippo}$ | $p_{Class}$ |
| --- | --- | --- | --- | --- | --- | --- | --- | --- | --- |
| SVM | PCA VF 99 | No | 66.78 % | 62.15 % | 79.56 % | 33.70 % |  |  | 0.621 |
| SVM | PCA VF 99 | Yes | 67.98 % | 63.14 % | 80.18 % | 35.30 % | 0.613 | 0.085 | 0.784 |
| SVM | PCA VF 95 | No | 66.72 % | 62.66% | 80.00 % | 34.30 % |  |  | 0.981 |
| SVM | PCA VF 95 | Yes | 67.82 % | 64.09 % | 79.71 % | 38.50 % | 0.660 | 0.086 | 0.628 |
| SVM | PCA VF 90 | No | 66.93 % | 63.65 % | 78.65 % | 39.10 % |  |  | 0.669 |
| SVM | PCA VF 90 | Yes | 67.58 % | 63.93 % | 78.67 % | 39.80 % | 0.806 | 0.064 | 0.356 |
| SVM | PCA VF 75 | No | 67.33 % | 64.99 % | 80.07 % | 40.30 % |  |  | 0.560 |
| SVM | PCA VF 75 | Yes | 69.29 % | 66.39 % | 80.82 % | 42.80 % | 0.371 | 0.169 | 0.879 |
| LR | PCA VF 99 | No | 65.42 % | 63.52 % | 87.46 % | 24.30 % |  |  |  |
| LR | PCA VF 99 | Yes | 67.35 % | 64.96 % | 87.76 % | 27.60 % | 0.513 | 0.059 |  |
| LR | PCA VF 95 | No | 66.79 % | 64.81 % | 88.34 % | 26.20 % |  |  |  |
| LR | PCA VF 95 | Yes | 68.77 % | 66.10 % | 86.07% | 33.30 % | 0.492 | 0.126 |  |
| LR | PCA VF 90 | No | 68.01 % | 64.95 % | 86.20 % | 30.10 % |  |  |  |
| LR | PCA VF 90 | Yes | 69.63 % | 65.08 % | 84.72 % | 32.90 % | 0.484 | 0.180 |  |
| LR | PCA VF 75 | No | 67.33 % | 68.22 % | 85.26 % | 36.50 % |  |  |  |
| LR | PCA VF 75 | Yes | 69.53 % | 66.20 % | 83.90 % | 37.20 % | 0.574 | 0.211 |  |

**Table 12:** Cross-validated performance measures with the non-QC dataset using AUC as the model selection criterion for different values of the RLR hyperparameter alpha testing high dimensional age removed features.

| Classifier | Feature Set | Alpha Value | AUC | ACC | SEN | SPE |
| --- | --- | --- | --- | --- | --- | --- |
| LR | Voxel based features | 0.25 | 71.06<br>$\pm 10.09$ | 62.88<br>$\pm 3.46$ | 95.28<br>$\pm 9.28$ | 9.80<br>$\pm 13.99$ |
| LR | Voxel based features | 0.5 | 71.34<br>$\pm 10.35$ | 66.99<br>$\pm 8.04$ | 82.68<br>$\pm 9.58$ | 41.30<br>$\pm 15.40$ |
| LR | Voxel based features | 0.75 | 72.37<br>$\pm 9.67$ | 66.65<br>$\pm 7.36$ | 86.70<br>$\pm 8.61$ | 33.80<br>$\pm 14.27$ |
| LR | Moradi features | 0.25 | 72.28<br>$\pm 9.28$ | 66.32<br>$\pm 6.35$ | 89.14<br>$\pm 11.18$ | 29.00<br>$\pm 27.51$ |
| LR | Moradi features | 0.5 | 74.04<br>$\pm 9.37$ | 70.84<br>$\pm 7.28$ | 86.79<br>$\pm 8.26$ | 44.70<br>$\pm 14.66$ |
| LR | Moradi features | 0.75 | 73.28<br>$\pm 9.11$ | 70.51<br>$\pm 6.65$ | 87.91<br>$\pm 9.75$ | 42.00<br>$\pm 16.25$ |
| LR | Region features | 0.25 | 80.18<br>$\pm 7.78$ | 71.72<br>$\pm 8.07$ | 83.45<br>$\pm 9.66$ | 52.50<br>$\pm 16.15$ |
| LR | Region features | 0.5 | 79.58<br>$\pm 7.71$ | 71.73<br>$\pm 7.56 \%$ | 84.07<br>$\pm 9.26$ | 51.50<br>$\pm 15.58$ |
| LR | Region features | 0.75 | 79.59<br>$\pm 7.54$ | 71.96<br>$\pm 7.97$ | 84.31<br>$\pm 9.63$ | 51.70<br>$\pm 15.24$ |

## Acknowledgments

Data collection and sharing for this project was funded by the Alzheimer’s Disease Neuroimaging Initiative (ADNI) (National Institutes of Health Grant U01 AG024904) and DOD ADNI (Department of Defense award number W81XWH-12-2-0012). ADNI is funded by the National Institute on Aging, the National In-stitute of Biomedical Imaging and Bioengineering, and through generous contributions from the following: AbbVie, Alzheimers Association; Alzheimers Drug Discovery Foundation; Araclon Biotech; BioClinica, Inc.; Biogen; Bristol-Myers Squibb Company; CereSpir, Inc.; Eisai Inc.; Elan Pharmaceuticals, Inc.; Eli Lilly and Company; EuroImmun; F. Hoffmann-La Roche Ltd and its affili-ated company Genentech, Inc.; Fujirebio; GE Healthcare; IXICO Ltd.; Janssen Alzheimer Immunotherapy Research & Development, LLC.; Johnson & Johnson Pharmaceutical Research & Development LLC.; Lumosity; Lundbeck; Merck & Co., Inc.; Meso Scale Diagnostics, LLC.; NeuroRx Research; Neurotrack Tech-nologies; Novartis Pharmaceuticals Corporation; Pfizer Inc.; Piramal Imaging; Servier; Takeda Pharmaceutical Company; and Transition Therapeutics.

J. Tohka’s work was supported by the Academy of Finland and V. Gómez-Verdejo’s work has been partly funded by the Spanish MINECO grant TEC2014-52289R, TEC2016-81900-REDT/AEI and TEC2017-83838-R.

## Footnotes

1 Information and data can be found at adni.loni.usc.edu

2 Available at adni.loni.usc.edu.

3 For up-to-date information, see www.adni-info.org.

4 http://surfer.nmr.mgh.harvard.edu/

5 Originally, this set included 274 measures. We selected a subset of 256 regions from the aforementioned 274 measures discarding the regions that presented missed data. A more detailed description of the 256 features is provided in https://github.com/MartaGomez/Regions-list-/wiki/Regions-list.

6 https://adni.bitbucket.io/reference/docs/UCSFFRESFR/UCSFFreeSurferMethodsSummary.pdf

7 http://scikit-learn.org/stable/modules/generated/sklearn.svm.SVC.html

8 https://www.csie.ntu.edu.tw/~cjlin/libsvm/

9 https://web.stanford.edu/~hastie/glmnet_python/>

10 Seehttps://github.com/MartaGomez/Regions-list-/wiki/Regions-list for a detailed description

